# Minor spliceosome disruption causes limb growth defects without altering patterning

**DOI:** 10.1101/2020.03.16.994384

**Authors:** Kyle D Drake, Christopher Lemoine, Gabriela S Aquino, Anna M Vaeth, Rahul N Kanadia

## Abstract

Disruption of the minor spliceosome causes primordial dwarfism in microcephalic osteodysplastic primordial dwarfism type 1. Similarly, primordial dwarfism in domesticated animals is linked to positive selection in minor spliceosome components. Despite the importance of minor intron splicing in limb size regulation, its role in limb development remains unexplored. Here we show that loss of U11 small nuclear RNA, an essential minor spliceosome component, results in stunted limbs that maintain patterning. Notably, earlier loss of U11 corresponded to increased severity. We find that limb size is reduced due to elevated minor intron retention in minor intron-containing genes that regulate cell cycle. Limb progenitor cells experience delayed prometaphase to metaphase transition and prolonged S-phase, resulting in death of rapidly dividing, distally located progenitors. Consequently, crucial limb patterning genes are upregulated and their expression is maintained spatially to achieve basic patterning. Overall, these findings reveal a potential mechanism shared in disease and domestication.

## Introduction

The minor spliceosome, which consists of five small nuclear RNAs (snRNAs) U11, U12, U4atac, U5, and U6atac and associated proteins, splices <0.5% of introns, termed minor introns, found in <2% of genes, termed minor intron-containing genes (MIGs)^1–3^. In metazoans, the minor spliceosome, MIGs, and the position of minor introns within MIGs are all highly conserved^4–6^. This conservation could be explained by the enrichment of MIGs in essential functions required for cell survival^7^, which is consistent with embryonic lethality observed in multiple animal models with constitutive loss of minor spliceosome function^8–10^. Consequently, it is unsurprising that human diseases with minor spliceosome loss-of-function mutations have not been discovered. However, mutations that result in partial loss of minor spliceosome function have been reported^11–16^.

For example, <u>m</u>icrocephalic <u>o</u>steodysplastic <u>p</u>rimordial <u>d</u>warfism type <u>1</u> (MOPD1), Roifman syndrome (RS), and Lowry-Wood syndrome (LWS) are linked to partial inhibition of the minor spliceosome^11,12,14,16^. Here, individuals harbor disparate mutations in *RNU4ATAC*, which encodes the U4atac snRNA, and display symptoms including microcephaly, micrognathia, vertebral deficits, and primordial dwarfism^16,17^. The severity of primordial dwarfism observed in MOPD1, RS, and LWS is observed along a gradation, with MOPD1 being the most severe, RS moderate, and LWS mild^16,17^. Nonetheless, in all cases, the basic patterning of the limb skeletal elements, including the presence of a stylopod (humerus; femur), zeugopod (radius/ulna; tibia/fibula), and autopod (hand; foot), is maintained^16,17^.

Since unique mutations in *RNU4ATAC* have been linked to MOPD1, RS, and LWS, the gradation of primordial dwarfism might be due to differential inhibition of the minor spliceosome^16,17^. This suggests that the activity of the minor spliceosome might serve as a rheostat for tissue size control, an idea bolstered by positive selection for genetic changes in MIGs and the minor spliceosome in domestication of several species^7^. In domestication, animals are selected for tameness, but secondarily show a constellation of phenotypic changes, referred to as domestication syndrome (DS)^18^. Symptoms of DS include neoteny, reduction in brain size, reductions in jaw and tooth size, changes in the number of vertebrae, floppy ears, curling of the tail, and reduction in limb size^18,19^. Notably, the phenotypes observed in DS parallel the symptoms observed in minor spliceosome-related diseases (Supp. Fig. 1)^7^. These observations led us to hypothesize that the activity of the minor spliceosome, and therefore MIG expression, plays a vital role in tissue size regulation. However, the underlying mechanism through which disruption of the minor spliceosome results in tissue size reduction while maintaining patterning remains unexplored.

Here we leverage our *Rnu11*^Flx/Flx^ mouse to inhibit the minor spliceosome in the developing limb by using *Prrx1*-Cre, which is expressed at embryonic day (E) 9.5 in the forelimb and E10.5 in the hindlimb^9,20^. We show that limb size is reduced upon minor spliceosome inhibition, such that the severity corresponded to the developmental time at which U11 was lost. Moreover, we show that the U11-null forelimb contained basic proximo-distal (shoulder to fingertip) patterning, whereas the U11-null hindlimb was patterned along the proximo-distal, antero-posterior (thumb to little finger), and dorso-ventral (back of hand to palm of hand) axes. We found that size reduction of the mutant limbs resulted from cell cycle defects and apoptosis of limb progenitor cells owing to elevated minor intron retention in MIGs crucial for cell cycle regulation. Moreover, our temporal RNAseq analysis showed a transcriptional response to the loss of U11 such that the expression of critical limb patterning genes, including *Shh, Fgf8, Hox* genes, and others, were upregulated to allow for proximo-distal patterning of the U11-null forelimb. In all, our study demonstrates potential cellular and molecular mechanisms underlying primordial dwarfism and micromelia in individuals with MOPD1, RS, and LWS^16,17^. Finally, our findings implicate how the activity of the minor spliceosome could be used to achieve tissue size reduction without loss of basic patterning in domestication.

## Results

### Differential effect of U11 loss on the developing forelimb and hindlimb

To test our hypothesis that inhibition of the minor spliceosome would lead to compromised limb growth, but not patterning, we generated *Rnu11^Flx/Flx^::Prrx1-*Cre^+^::*CAG*-loxpSTOPloxp-*tdTomato^+^* (mutant) embryos that we compared to wild-type (WT) littermates. We used tdTomato expression as a reporter of Cre activity to corroborate the previously described expression difference of *Prrx1* in the forelimb and hindlimb buds^20^. As expected, we found tdTomato signal only in the forelimb at E10.5 but in both the forelimb and hindlimb at E11.5 (Supp. Fig. 2a-b). Moreover, by whole mount *in* situ hybridization (WISH), we confirmed the loss of U11 snRNA in the E10.5 mutant forelimb, but not hindlimb (Fig. 1a). Furthermore, quantitative reverse transcription-PCR (qRT-PCR) revealed a significant, 62.9% reduction of U11 in the E10.5 mutant forelimb, whereas the hindlimb did not show similar loss (Fig. 1b). At E11.5, qRT-PCR for U11 showed that it was significantly downregulated 75.1% in the mutant forelimb and 68.8% in the mutant hindlimb (Fig. 1b). This temporal difference in U11 loss was reflected in the early onset of visible morphological defects in the mutant forelimb at E11.5, with increased severity observed at E12.5 and E13 (Fig. 1c; Supp. Fig. 3a). Quantification of limb bud surface area confirmed that the E11.5 mutant forelimb was significantly smaller than the E11.5 WT forelimb, and that this phenotype was exacerbated by E12.5 (Fig. 1d). While, the mutant hindlimb did not show gross morphological defects by E13, quantification revealed that the overall size of the mutant hindlimb was significantly reduced by E12.5 (Fig. 1c-d; Supp. Fig. 3a). There was no left/right bias observed in the mutant forelimb or hindlimb phenotypes (Supp. Fig. 3b).

**Figure 1.**
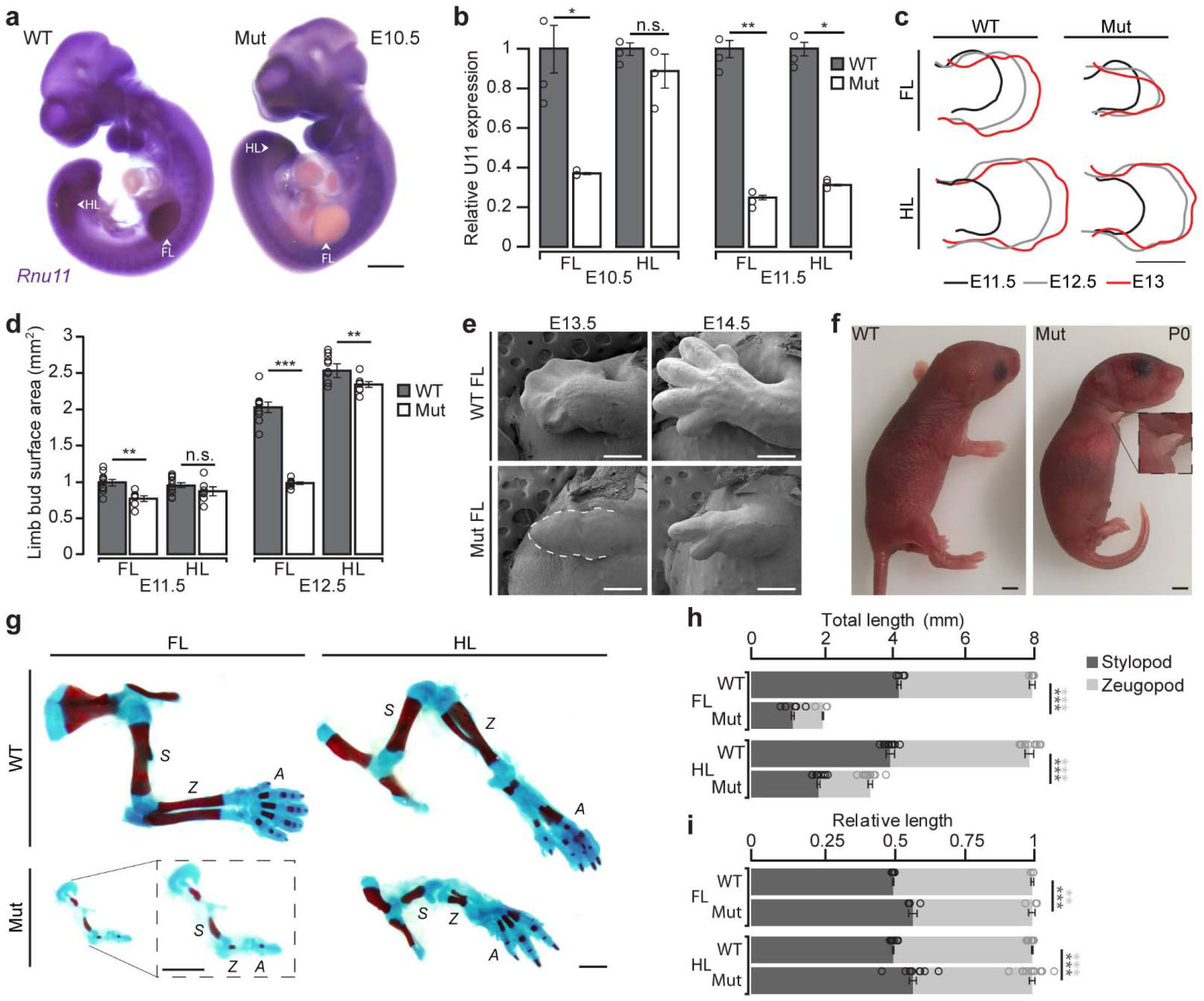
U11-null limbs have stunted growth but maintain basic patterning. **(a)** Whole mount *in situ* hybridization (WISH) for *Rnu11* in E10.5 WT and mutant (mut) embryos with arrows pointing to forelimb (FL) and hindlimb (HL) buds. **(b)** Quantification of WT and mutant *Rnu11* expression from dissected (n=3) E10.5 and E11.5 FL and HL buds through qRT-PCR. **(c-d)** Limb bud traces from WT and mut FL and HL at E11.5 (black), E12.5 (grey), and E13 (red) (**c**) with quantification of surface area **(d)**. **(e)** SEM images of WT and mutant FL buds at E13.5 and E14.5. **(f)** P0 images of WT and mut pups with higher magnification image of mut FL. **(g)** Skeletal preparation for WT and mut FL and HL at P0 with higher magnification image of mut FL. **(h-i)** Quantification of total **(h)** and relative **(i)** long bone length for WT and mut FL and HL at P0. S=stylopod, Z=zeugopod, A=autopod. Scale bars represent 100 um in **(a)**, **(c)**, **(e)**, **(g)**; 1mm in **(f)**. All bar charts represent mean with error bars representing standard error of the mean. Significance determined via student’s two-tailed T-test. *=*p*<0.05, **=*p*<0.01, ***=*p*<0.001.

### U11-null limbs are reduced in size but show basic patterning at birth

Based on the reduced size of the mutant limb buds, we employed scanning electron micrograph (SEM) imaging at E13.5 and E14.5 to determine whether patterning was also affected. At E13.5, the mutant forelimb did not appear to have an antero-posterior axis as it lacked digital condensations (Fig. 1e). However, at E14.5, the mutant forelimb showed two digital processes, indicating the presence of a proximo-distal and antero-posterior axis (Fig. 1e). The mutant hindlimb did not show distinct morphological abnormalities at either time point imaged (Supp. Fig. 3c). The presence of digits in the E14.5 mutant forelimb led us to hypothesize that the other elements of the proximo-distal axis, including the stylopod (humerus) and zeugopod (radius/ulna) were also formed. To test this hypothesis, we performed skeletal preparations on WT and mutant mice at birth (P0), which showed that, although severely stunted, the P0 mutant forelimb contained three distinct skeletal units: a humerus, a single zeugopod element, and a single digit (Fig. 1f-h). Moreover, the mutant hindlimb, which was smaller compared to the WT, consisted of a femur, tibia, fibula, and foot (Fig. 1f-h). While the mutant hindlimb lacked a medial cuneiform and had a stunted digit 1, basic patterning across all three axes was preserved (Fig. 1f-g; Supp. Fig. 4a).

Given the reduction in overall limb size, we aimed to determine whether the proximal and distal long bones of the mutant limbs were proportionally reduced in size. We found that the stylopod-zeugopod ratio was significantly altered in the mutant forelimb and hindlimb (Fig. 1i). In both, the stylopod comprised a significantly greater amount of total limb length, indicating that the mutant limbs were mesomelic (Fig. 1i). Furthermore, in MOPD1, osteogenic defects causing narrow bones with reduced bone density have been reported^21^. Therefore, we quantified the length and width of the Alizarin red-stained (ossified) region of the long bones of the WT and mutant limbs at P0. We discovered that the stylopod and zeugopod were significantly narrower and less ossified in both the mutant forelimb and hindlimb relative to WT controls (Supp. Fig. 4b-d).

### U11 loss results in elevated minor intron retention in MIGs that regulate cell cycle

To identify the underlying molecular pathways disrupted in the mutant limbs, we performed total RNAseq using E10.5 and E11.5 WT and mutant forelimb and hindlimb samples. We found that U11 was reduced by 73.5% at E10.5 and 81.2% at E11.5 in the mutant forelimb compared to their respective WT controls (Fig. 2a). Similarly, we found that U11 was reduced by 62.5% at E10.5 and 75.8% at E11.5 in the mutant hindlimb compared to their respective WT controls (Fig. 2a). The incomplete loss of U11 expression is consistent with *Prrx1*-Cre being expressed in the limb bud mesoderm but not in the overlying limb bud ectoderm^20^. Moreover, forelimb and hindlimb specificity was reflected in the significant enrichment of *Tbx5*, a forelimb marker, in all forelimb samples, significant enrichment of *Tbx4*, a hindlimb marker, in all hindlimb samples, and non-differential expression (NonDE) of *Prrx1* in all sample comparisons (Fig. 2b)^22^.

**Figure 2.**
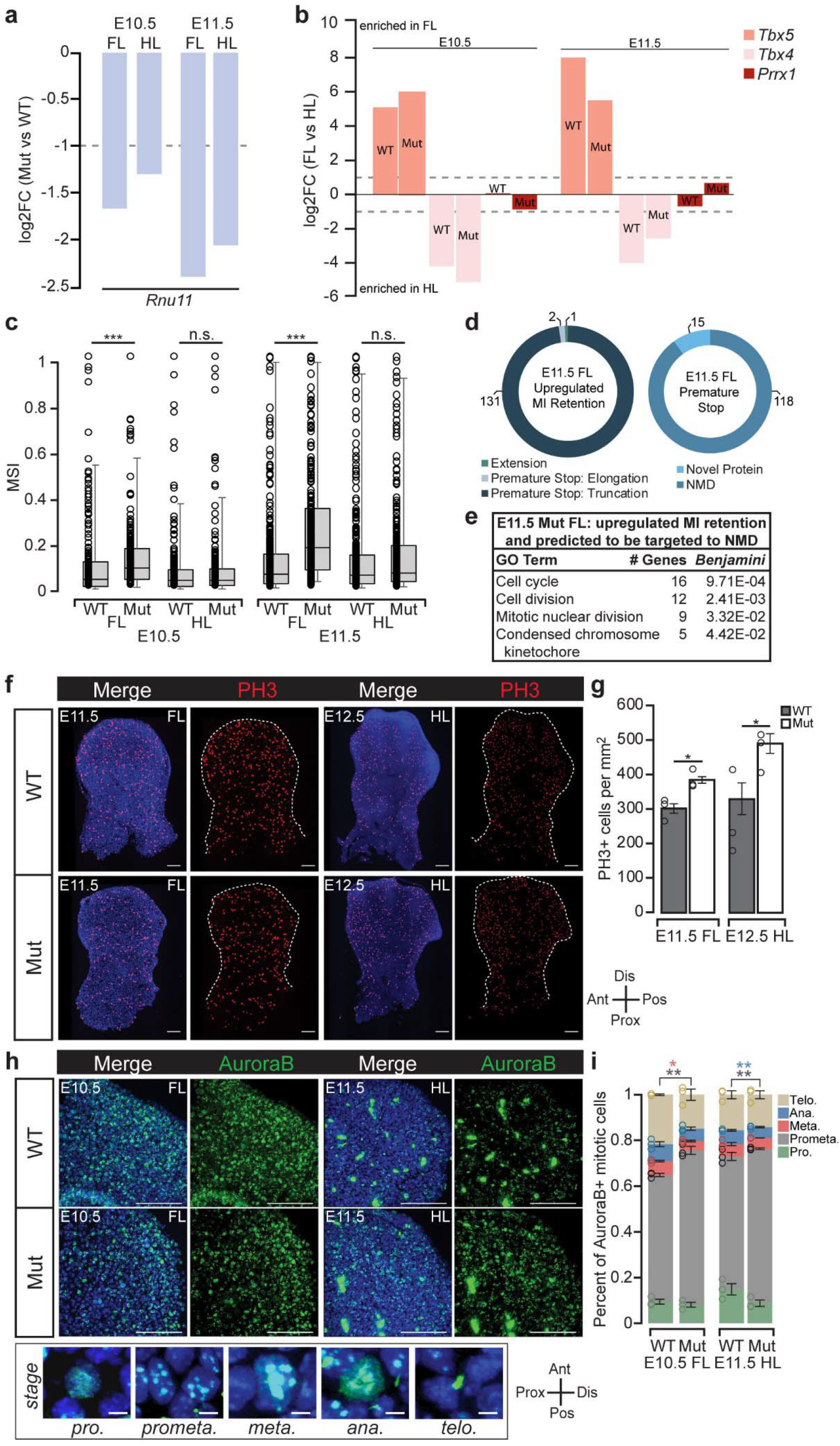
U11 loss results in minor intron retention in MIGs that regulate cell cycle and causes defective mitosis. **(a)** Bar chart of confident log2 fold change (FC) of *Rnu11* in the mutant (mut) forelimb (FL) and hindlimb (HL) (compared to WT age- and tissue-matched controls) at E10.5 and E11.5 quantified through RNAseq with dashed line representing threshold for significance. **(b)** Bar chart of confident log2FC of *Tbx5*, *Tbx4*, and *Prrx1* in WT and mutant FL and HL at E10.5 and E11.5 quantified through RNAseq with dashed lines representing thresholds for significance. **(c)** Box-plot representing 10^th^-90^th^ percentile mis-splicing index (MSI) for all MIGs that show minor intron retention in WT and mutant FL and HL at E10.5 and E11.5. **(d)** Pie chart depicting consequence of minor intron retention on the open reading frame of MIGs with significantly elevated minor intron retention in the E11.5 mutant FL (left) and whether a premature stop codon is predicted to trigger nonsense mediated decay (NMD) or result in novel protein production (right). **(e)** GO enrichment for MIGs that have significantly upregulated minor intron (MI) retention and are predicted to be targeted to NMD in the E11.5 mutant FL. **(f-g)** Immunofluorescence (IF) for phospho-histone H3 (PH3) counterstained with DAPI in WT and mutant E11.5 FL and E12.5 HL **(f)** with quantification **(g)** normalized to limb bud area. **(h-i)** IF for AuroraB counterstained with DAPI in WT and mutant E10.5 FL and E11.5 HL **(h)** with quantification **(i)** of the percent of cells in each mitotic stage per sample. Ant=anterior, Post=posterior, Dis=Distal, Prox=Proximal. Scale bars represent 100 um. Bar charts in **(g)**, **(i)** represent mean and error bars represent standard error of the mean. Significance determined by Kruskal-Wallis test **(c)** and student’s two-tailed T-test in **(g)**, **(i)**. *=*p*<0.05, **=*p*<0.01, ***=*p*<0.001.

Since U11 loss is expected to inhibit the minor spliceosome, we performed minor intron retention analysis on our RNAseq data. We discovered elevated minor intron retention in the E10.5 mutant forelimb compared to its WT counterpart as indicated by a significantly elevated median mis-splicing index (MSI) (Fig. 2c). In contrast, there was no significant difference in minor intron retention between the E10.5 WT and mutant hindlimb (Fig. 2c). Similarly, at E11.5, we found a significantly elevated median MSI in the mutant forelimb but not in the mutant hindlimb (Fig. 2c). For each sample, we next identified the individual minor introns that were retained at significantly higher levels compared to the corresponding WT data set. We found 21 minor introns to be significantly retained in the E10.5 mutant forelimb, whereas only two minor introns showed significantly elevated retention in the E10.5 mutant hindlimb compared to their respective WT controls (Supp. Data File 1). At E11.5, we found 134 minor introns to be significantly retained in the mutant forelimb and 15 minor introns to be significantly retained in the mutant hindlimb compared to their respective WT controls (Supp. Data File 1). Altogether, retention of these minor introns affected splicing of 152 MIGs in at least one of the mutant conditions.

Since minor intron retention can introduce a premature stop codon and result in nuclear degradation or nonsense mediated decay (NMD), we analyzed the effect of minor intron retention on the open reading frame (ORF) of these 152 MIGs^3,9^. Indeed, we found that minor intron retention almost invariably resulted in the introduction of a premature stop codon (150/152; 98.6%) (Supp. Data File 2; Supp. Table 1). Of these, we found that 134 events were predicted to result in NMD, whereas only 16 would encode novel proteins (Supp. Table 1). We next used DAVID to identify the pathways that may be compromised upon loss-of-function of these MIGs. Only the 118 MIGs in the E11.5 mutant forelimb predicted to undergo NMD had significant enrichment, including cell cycle, cell division, mitotic nuclear division, and condensed chromosome kinetochore (Fig. 2d-e; Supp. Data File 3). Thus, this suggested potential cell cycle defects in U11-null limbs.

### U11 loss results in defective mitosis, prolonged S-phase, and slowing of cell cycle

To test whether cell cycle was affected in the E11.5 mutant forelimb, we first performed immunofluorescence (IF) for phospho-histone H3 (PH3), which marks all mitotic cells. Indeed, we observed an increase in PH3+ cells at E11.5 in the mutant forelimb, but not at E10.5 (Fig. 2f-g; Supp. Fig. 5a-b). Similarly, we observed an increase in the number of PH3+ cells in the mutant hindlimb at E12.5, but not at E11.5 (Fig. 2f-g, Supp. Fig. 5a, 5c). The increase in mitotic cells in the E11.5 mutant forelimb and E12.5 mutant hindlimb could be caused by a delay in progenitor cell progression through the mitotic phases, as we previously found in the U11-null pallium^9^. Therefore, we performed IF for AuroraB, as its subcellular localization can be used to identify prophase, prometaphase, metaphase, anaphase, and telophase (Fig. 2h). While the total number of cells in mitosis was not different between the E10.5 WT and mutant forelimb, the distribution of cells within each phase was shifted such that there were significantly more cells in prometaphase and significantly fewer cells in metaphase in the mutant (Fig. 2h-i). Similarly, in the E11.5 mutant hindlimb, we found significantly more cells in prometaphase and significantly fewer cells in anaphase (Fig. 2h-i). Taken together, these data suggested that U11 loss caused defective mitotic progression and an accumulation of PH3+ cells 24 hours later, indicating that cell cycle speed might be generally reduced in limb progenitor cells.

To determine the cell cycle speed of limb progenitor cells, we employed a BrdU/EdU pulse-chase paradigm previously used in the developing limb by Boehm et al.^23^ (Fig. 3a). According to this paradigm, pregnant dams were injected with BrdU 2 hours before harvest so that all cells in S-phase within 30 minutes of injection were marked (Fig. 3a)^23^. Subsequently, EdU injection 30 minutes prior to harvest marked all cells in S-phase at the time of harvest (Fig. 3a)^23^. This double-labeling technique allowed for the identification of cells that did not enter S-phase in the two-hour period, stayed in S-phase during the two-hour period, or left S-phase prior to the EdU injection (Fig. 3a)^23^. In turn, quantification of these populations of cells allowed for the estimation of S-phase length and total cell cycle time^23^. Using this strategy, we discovered a significant increase in the length of S-phase and total cell cycle time in the E10.5 mutant forelimb compared to its WT counterpart (Fig. 3b-c). Specifically, S-phase was increased by 3.29 hours, whereas cell cycle time was increased by 4.08 hours (Fig. 3c). Similarly, we found a significant increase in S-phase by 5.88 hours and cell cycle time by 6.81 hours in the E11.5 mutant hindlimb relative to its WT counterpart (Fig. 3b, 3d). Thus, in both the E10.5 mutant forelimb and E11.5 mutant hindlimb, mitotic delay and prolonged S-phase contribute to decreased cell cycle speed, but the lengthening of S-phase was the primary driver (Fig. 2i, 3c-d). Defective cell cycle progression and subsequently reduced cell cycle speed led us to hypothesize that rapidly dividing cells would undergo apoptosis, as we have previously observed^9^.

**Figure 3.**
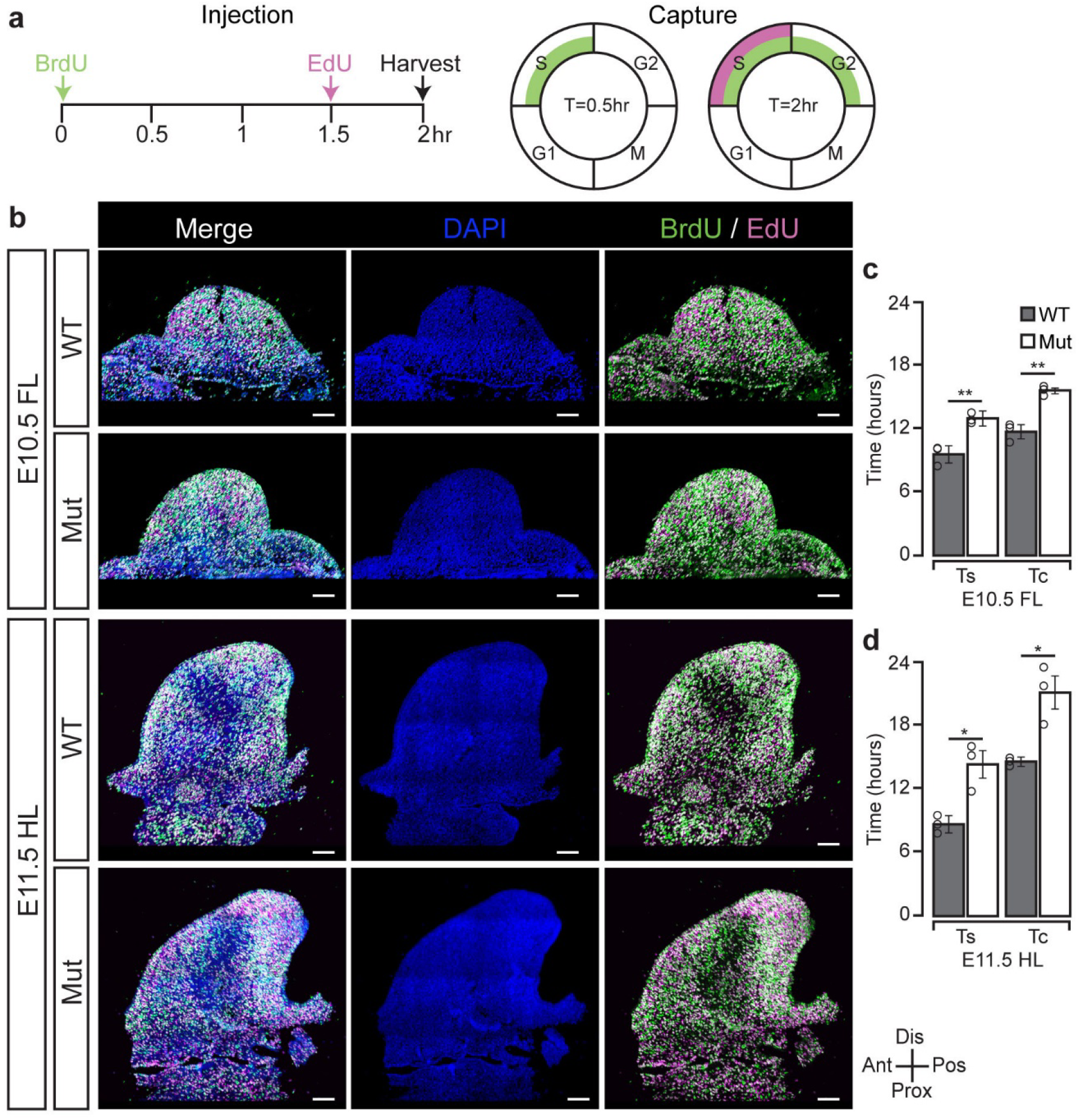
U11 loss results in an elongation of S-phase and a reduction of cell cycle speed. **(a)** Adapted schematic of injection paradigm to quantify cell cycle speed as described by Boehm et al.,^23^. BrdU and EdU take 30 minutes to be detected, and are thus captured at T=0.5 hours (hr) and T=2hr, respectively. **(b)** IF for BrdU and enzymatic detection for EdU with DAPI counterstain in WT and mutant (mut) E10.5 forelimb (FL) and E11.5 hindlimb (HL). **(c)** Quantification of total S-phase time (Ts) and total cell cycle time (Tc) for WT and mutant E10.5 FL and E11.5 HL. Ant=anterior, Post=posterior, Dis=Distal, Prox=Proximal. Bar charts represent mean and error bars represent standard error of the mean. Scale bars represent 50 um. Significance determined by student’s two-tailed T-test. *=*p*<0.05, **=*p*<0.01.

### Loss of U11 results in death of a subset of distally concentrated limb progenitor cells

To identify the U11-null limb progenitor cells that undergo apoptosis, we performed fluorescence *in situ* hybridization (FISH) for U11 followed by terminal dUTP nick end labeling (TUNEL) to mark dying cells. Across the dorso-ventral axis, different sections of the limb showed varying patterns of TUNEL. Therefore, we sought to obtain a 3-dimensional perspective of apoptosis by processing all sections of the developing limb followed by ImageJ to reconstruct a 2D heat map for TUNEL. Using this strategy, we found mesodermal U11 loss with a significant increase in cell death concentrated distally and posteriorly in the E11.5 mutant forelimb, but not in the E10.5 mutant forelimb (Fig. 4a, 4c-e; Supp. Fig. 6a-b). Similarly, the mutant hindlimb showed significant cell death concentrated distally and anteriorly at E12.5, but not at E11.5 (Fig. 4b-e; Supp. Fig. 6a, 6c). Thus, cell cycle defects caused by elevated minor intron retention led to apoptosis in the mutant limbs (Fig. 2c, 2e, 2i, 3c-d, 4c). Despite these defects, basic proximo-distal patterning and complete patterning were observed in the P0 mutant forelimb and hindlimb, respectively (Fig. 1g). Therefore, we hypothesized that the developing mutant limbs undergo transcriptome alterations to maintain the expression of genes crucial for limb patterning.

**Figure 4.**
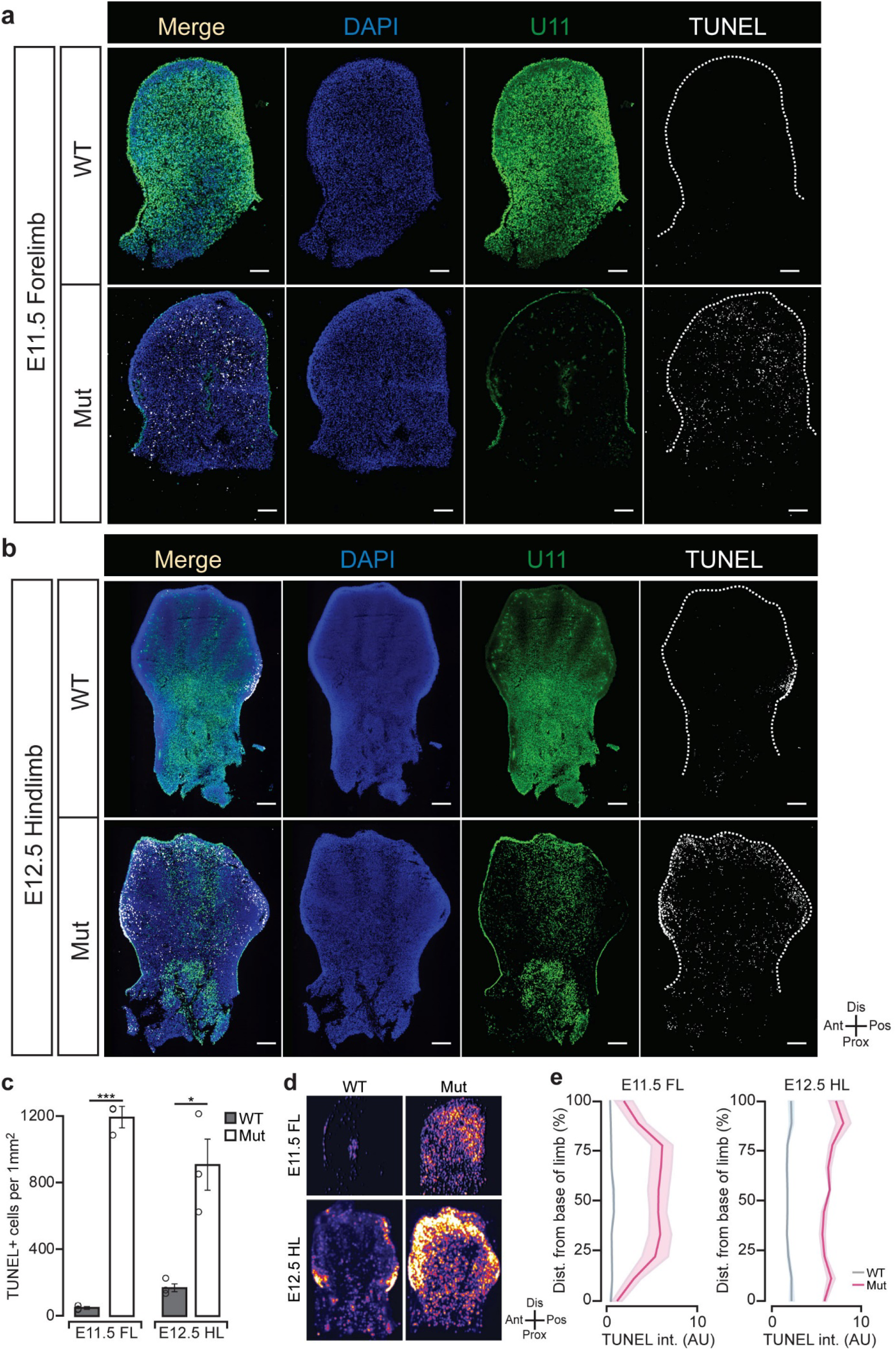
U11 loss causes distally concentrated cell death. **(a-b)** Fluorescent *in situ* hybridization (FISH) for *Rnu11* with terminal dUTP-nick end labeling (TUNEL) counterstained with DAPI in WT and mutant (mut) E11.5 forelimb (FL) **(a)** and E12.5 hindlimb (HL) **(b)**. **(c)** Quantification of TUNEL+ cells normalized to limb bud area. **(d)** Heat map of serially overlaid images from **(a-b)** with brighter color representing higher concentration of TUNEL+ signal. **(e)** Quantification of TUNEL intensity from **(d)** in arbitrary units (AU) in relation to the distance (dist.) from the base of the limb bud. Ant=anterior, Post=posterior, Dis=Distal, Prox=Proximal. Bar charts represent mean and error bars in **(c)** and shaded region in **(e)** represent standard error of mean. Scale bars represent 100 um. Significance determined via student’s two-tailed T-test. *=*p*<0.05, **=*p*<0.01,***=*p*<0.001.

### RNAseq analysis reveals upregulation of limb patterning genes in the E11.5 mutant forelimb

To determine whether transcriptomic alterations were occurring in the mutant limb buds, we examined the expression of all protein-coding genes. We considered a gene expressed if it had transcripts per million (TPM) above 0.50, the lowest expression of *Shh*, a key limb patterning gene, found among the two E10.5 WT data sets. With this threshold, we performed pairwise comparisons for each age- and tissue-matched sample. We found that 62 genes were upregulated (≥2 fold change (FC); *p*≤0.01) and 245 genes were downregulated (≤2 FC; *p*≤0.01) in the E10.5 mutant forelimb compared its WT counterpart (Fig. 5a; Supp. Data File 4). In the E10.5 mutant hindlimb, 145 genes were upregulated and 97 genes were downregulated (Fig. 5a; Supp. Data File 4). At E11.5, 1,004 genes were upregulated and 16 genes were downregulated in the mutant forelimb (Fig. 5a; Supp. Data File 4). Lastly, 7 genes were upregulated and 65 genes were downregulated in the E11.5 mutant hindlimb (Fig. 5a; Supp. Data File 4). We next leveraged DAVID to extract the biological pathways that might be affected by the differentially expressed gene sets. While there were GO terms enriched by genes differentially expressed in all pairwise comparisons, only the 1,004 upregulated genes in the E11.5 mutant forelimb enriched for limb patterning-related GO terms (Supp. Data File 3; Supp. Table 2). Specifically, these genes enriched for 101 GO pathways, including anterior/posterior pattern specification, embryonic skeletal system morphogenesis, and limb morphogenesis (Fig. 5b; Supp. Data File 3). Moreover, we discovered enrichment in specific pathways known to be compromised in the mutant limb, including U12-type (minor) spliceosomal complex, cell cycle, and cell death (Fig. 5b; Supp. Data File 3).

**Figure 5.**
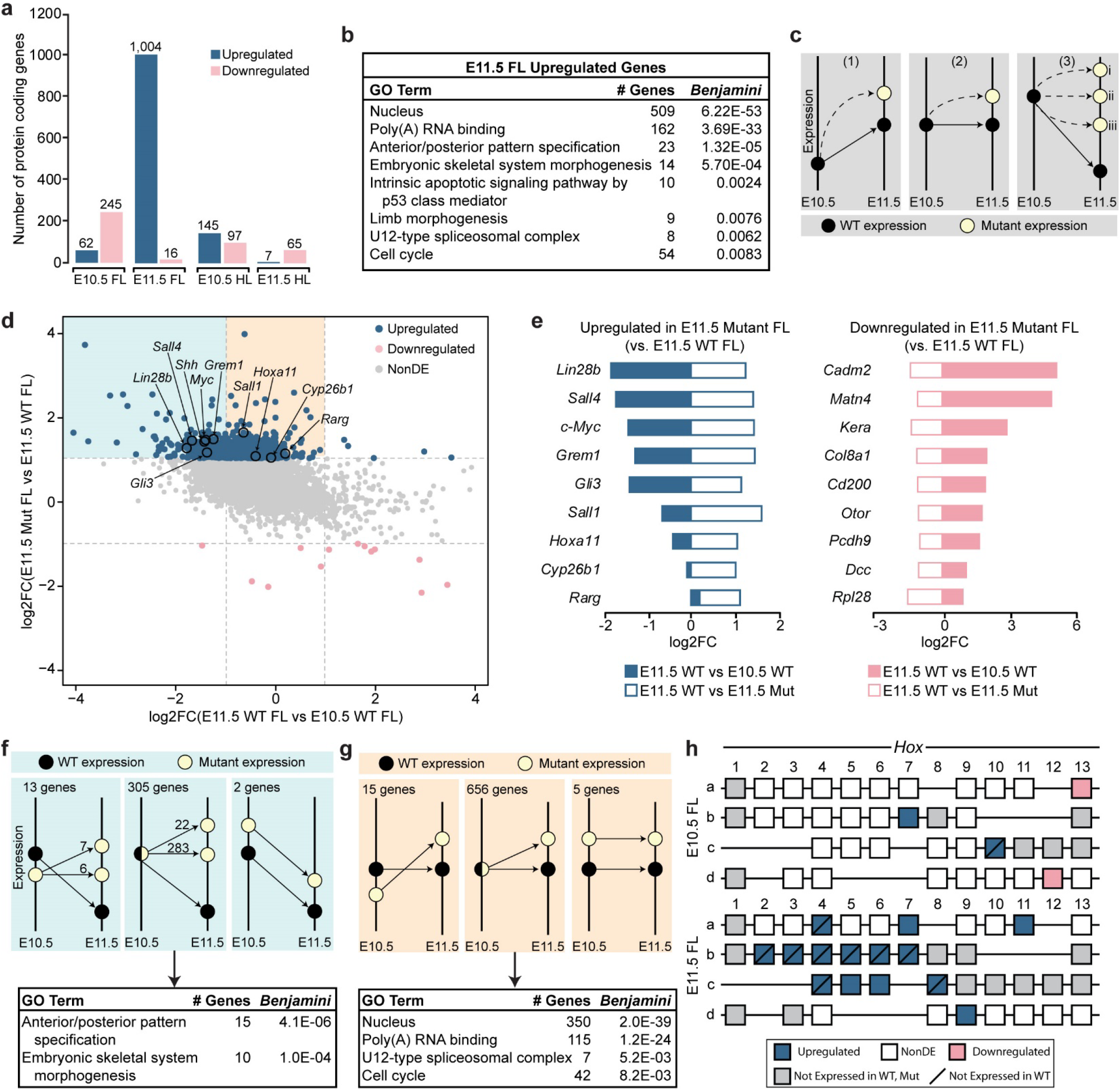
Loss of U11 results in prevented downregulation of critical limb patterning genes in the E11.5 mutant forelimb. **(a)** Bar chart representing the number of upregulated and downregulated protein-coding genes in the mutant (mut) E10.5 and E11.5 forelimb (FL) and hindlimb (HL) (when compared to their age- and tissue-matched WT controls). **(b)** GO pathway enrichment for upregulated genes in the E11.5 mutant FL (when compared to the E11.5 WT FL). **(c)** Schematic depicting three possible manners of gene upregulation in the E11.5 mutant (yellow) FL relative to that gene’s E10.5 WT (black) expression level. **(d)** Cumulative plot showing the confident log2 fold-change (log2FC) of all expressed protein coding genes in the E10.5 WT FL, E11.5 WT FL, and E11.5 mutant FL with color denoting differential expression in the E11.5 WT FL vs E11.5 mutant FL comparison. **(e)** Bar chart showing the confident log2FC of upregulated (left) and downregulated (right) genes in the E11.5 mutant FL (compared to the E11.5 WT FL) relative to WT FL temporal expression. **(f-g)** Schematic depicting the number of genes that are downregulated **(f)** or transcriptionally stable **(g)** from E10.5 to E11.5 in WT FL but are upregulated in the E11.5 mutant FL (when compared to the E11.5 WT FL) with inclusion of the transcriptional change from the E10.5 mutant FL to the E11.5 mutant FL. DAVID analysis on the highest represented group in **(f)** and **(g)** reported below. **(h)** Schematic depicting transcriptional changes of all *Hox* genes in the E10.5 and E11.5 mutant forelimb (as compared to their WT age- and tissue-matched controls). NonDE=non-differentially expressed.

### Genes involved in limb patterning show E10.5 WT-like expression in the E11.5 mutant forelimb

The enrichment of limb morphogenesis pathways by the 1,004 upregulated genes in the E11.5 mutant forelimb led us to hypothesize that the mutant transcriptome was attempting to maintain basic patterning (Fig. 5b). To test this hypothesis, we determined whether these genes were actively upregulated or were prevented from being downregulated. Therefore, we examined the changes in expression of these genes from E10.5 to E11.5 in the WT forelimb. In this comparison, a gene can either be upregulated, not changed, or downregulated (Fig. 5c). If a gene is upregulated from E10.5 to E11.5 in the WT forelimb and was found to be upregulated in the E11.5 mutant forelimb (compared to the E11.5 WT), it must have been actively upregulated in the mutant (Fig. 5c, panel 1). Alternatively, a gene may remain unchanged in its expression from E10.5 to E11.5 in the WT forelimb. If that gene is upregulated in the E11.5 mutant forelimb compared to the E11.5 WT forelimb, it would also be considered actively upregulated (Fig. 5c, panel 2). Lastly, a gene may downregulated from E10.5 to E11.5 in the WT forelimb. For this gene to be upregulated in the E11.5 mutant forelimb, it could have either (i) been actively upregulated, (ii) maintained its E10.5 level of expression, and thus was prevented from being downregulated, or (iii) undergone a decrease in expression but not to the extent it normally would (Fig. 5c, panel 3). To assess these possibilities, we isolated all protein-coding genes expressed in the E10.5 WT, E11.5 WT, and E11.5 mutant forelimb data sets. We then plotted the log2FC of each gene in the E10.5 WT vs E11.5 WT forelimb comparison (x-axis), which represents the normal change in gene expression during limb development, against the log2FC of each gene in the E11.5 WT vs E11.5 mutant forelimb comparison (y-axis), which represents the changes in gene expression in the E11.5 mutant forelimb (Fig. 5d). This cumulative plot indicated that most upregulated genes in the E11.5 mutant forelimb fall into two categories: (1) genes that are transcriptionally downregulated between E10.5 and E11.5 in the WT forelimb, such as *Lin28b*, *Gli3, Sall4, Shh, c-Myc,* and *Grem1* (light blue; 320 genes; Fig. 5c, panel 3); or (2) genes that are transcriptionally maintained from E10.5 to E11.5 in the WT forelimb but show an increased expression in the mutant, such as *Sall1, Hoxa11, Cyp26b1,* and *Rarg* (beige; 676 genes; Fig. 5c, panel 2) (Fig. 5d-e). In contrast, of the genes that were downregulated in the E11.5 mutant forelimb compared to the E11.5 WT forelimb, most were found to be upregulated in the E11.5 WT forelimb compared to the E10.5 WT forelimb (Fig. 5d-e).

The changes we observed in the E11.5 mutant forelimb suggested that it was transcriptionally responding to cell cycle defects and apoptosis precipitating between E10.5 and E11.5. To determine this, we broke down the two primary sets of upregulated genes into whether they were upregulated, NonDE, or downregulated, when comparing the E10.5 WT forelimb with the E10.5 mutant forelimb (Fig. 5f-g). Of the 320 genes that were upregulated in the E11.5 mutant forelimb (compared to the E11.5 WT forelimb) when they are normally downregulated in the E11.5 WT forelimb (compared to the E10.5 WT forelimb) (Fig. 5f), 13 were downregulated in the E10.5 mutant forelimb compared to the E10.5 WT forelimb (Fig. 5f, panel 1), 305 were NonDE in this comparison (Fig. 5f, panel 2), and 2 were upregulated in the E10.5 mutant forelimb (Fig. 5f, panel 3). Moreover, of the 676 genes that were transcriptionally maintained from E10.5 to E11.5 in the WT forelimb but show an increased expression in the E11.5 mutant forelimb (compared to the E11.5 WT forelimb) (Fig. 5g), 15 were downregulated in the E10.5 mutant forelimb compared to the E10.5 WT forelimb (Fig. 5g, panel 1), 656 were NonDE in this comparison (Fig. 5g, panel 2), and 5 were upregulated in the E10.5 mutant forelimb (Fig. 5g, panel 3). This analysis revealed that a majority of the significantly upregulated genes in the E11.5 mutant forelimb were NonDE in the E10.5 mutant forelimb compared to the E10.5 WT forelimb (Fig. 5f, panel 2, 5g, panel 2).

From this analysis, we were able to refine our understanding of the temporal kinetics of the genes that previously enriched for the GO terms such as anterior/posterior pattern specification and embryonic skeletal system morphogenesis (Fig. 5b). Indeed, these same two GO terms are enriched when we performed DAVID analysis on the 283 genes that were prevented from being downregulated in the E11.5 mutant forelimb (Fig. 5f, panel 2; Supp. Data File 3). Similarly, the previous GO terms U12-type (minor) spliceosomal complex and cell cycle were found to be generated again when we performed DAVID analysis on the 656 genes that were actively upregulated in the E11.5 mutant forelimb when they should have stable temporal expression (Fig. 5b, 5g, panel 2; Supp. Data File 3).

Moreover, since 10 out of the 15 genes which enriched for anterior/posterior patterning were *Hox* genes, and it is known that these genes also play a critical role in proximo-distal patterning^24^, we compared the expression of the entire *Hox* gene cluster between the WT and mutant forelimb at E10.5 and E11.5. We found that, at E10.5, *Hoxb7* and *Hoxc10* were upregulated whereas *Hoxa13* and *Hoxd12* were downregulated in the mutant (Fig. 5h). In contrast, at E11.5, *Hoxa4, a7, a11, Hoxb2, b3, b4, b5, b6, b7, Hoxc4, c5, c6, c8,* and *Hoxd9* were upregulated, while none were downregulated in the mutant (Fig. 5h). These findings demonstrate significant transcriptomic upregulation of early limb patterning genes^24^.

### The spatial expression of limb patterning genes is maintained in the E11.5 mutant forelimb

Our temporal RNAseq analysis showed that crucial limb patterning genes were upregulated in the E11.5 mutant forelimb (Fig. 5b). Moreover, the E11.5 mutant forelimb was found to have cell death concentrated distally and posteriorly, which could implicate gene expression of cells in these regions (Fig. 4d). Thus, we wanted to assess whether these changes in gene expression were maintained spatially, as this dimension of expression is crucial for limb patterning. We selected *Shh*, which is crucial for establishing both proximo-distal and antero-posterior patterning; *Hoxa11*, as it is expressed by zeugopod progenitors; *Hoxa13*, as it is expressed by autopod progenitors; *Sall1,* due to its interaction with *Shh* and *Hoxa13;* and *Fgf8* and *Cyp26b1*, for their crucial roles in proximo-distal patterning^22,25,26^. WISH analysis showed that *Shh, Sall1, Hoxa11, Hoxa13, Fgf8,* and *Cyp26b1* possess expression domains in the E11.5 mutant forelimb that are comparable to the E11.5 WT forelimb (Fig. 6a-b). Thus, through RNAseq and WISH, we found that U11 loss resulted in a temporal transcriptome change to maintain spatial expression domains of limb patterning genes, thereby allowing the basic patterning of the mutant forelimb.

**Figure 6.**
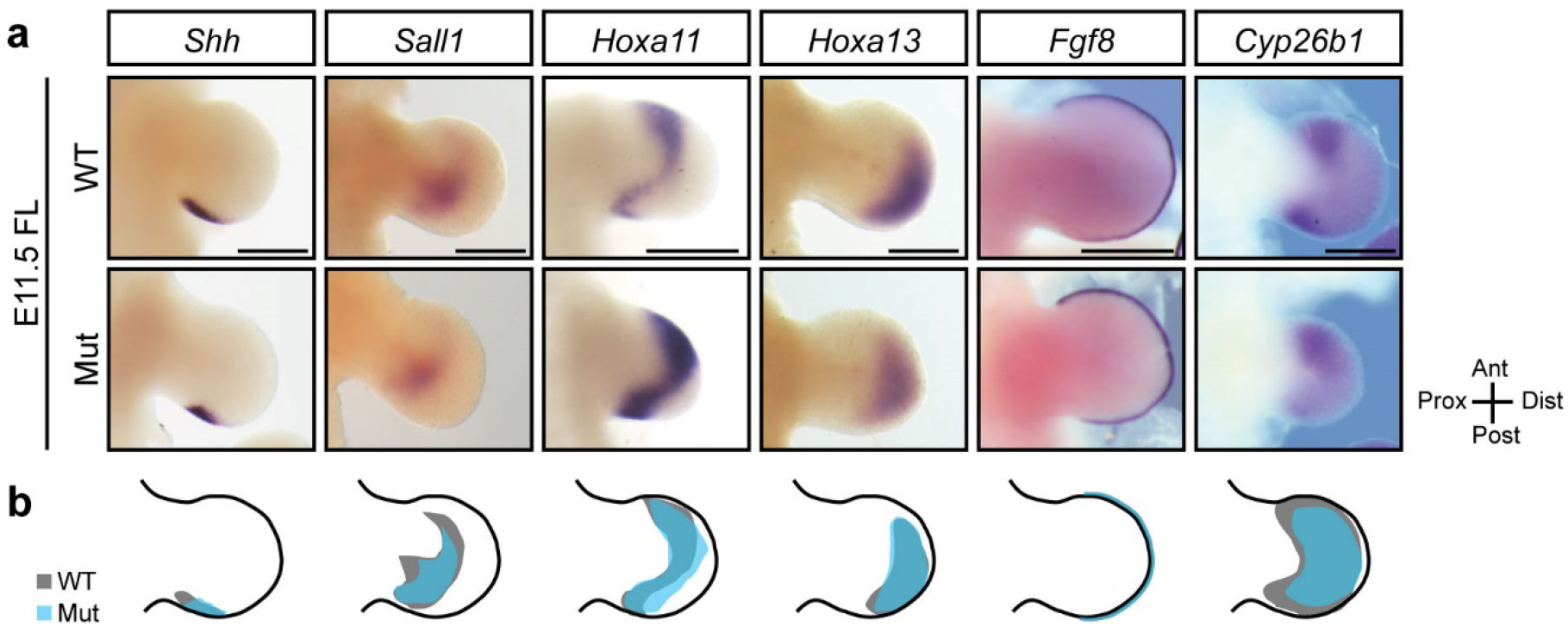
U11 loss does not alter the expression domains of limb patterning genes in the E11.5 mutant forelimb. **(a)** WISH for *Shh, Sall1, Hoxa11, Hoxa13, Fgf8*, and *Cyp26b1* in the WT and mutant (mut) E11.5 forelimb (FL). **(b)** Schematic showing traced WT limb bud and gene expression domain (black) with mutant gene expression domain (blue) overlaid. Scale bars represent 50 um.

## Discussion

The regulation of tissue size is a fundamental question in biology as, in mammals, the adult body mass ranges from ~1.5 grams in the Etruscan shrew to ~1.5 million grams in the blue whale^27^. A direct example of tissue size reduction is observed in the limb size of domesticated animals, such as dogs, cats, and cows, compared to their wild ancestors^19^. Comparisons of genome-wide association studies between domesticated animals and their wild counterparts have revealed shared selection for genetic changes in MIGs and components of the minor spliceosome^7^. This finding suggests that tissue size could be altered by modulating expression of certain MIGs and/or the minor spliceosome, which is reinforced by the symptomology of MOPD1, RS, and LWS^7^. An important characteristic of limb size reduction in domestication syndrome and minor spliceosome-related disease is the maintenance of limb patterning^16,17,19^. In agreement, we show here that U11-null limbs are reduced in size but maintain patterning at birth (Fig. 1g-h). In the mutant forelimb, the discordance between our observation of a single digit at P0 and two digits through SEM at E14.5 is most likely due to the penetrance of *Prrx1*-Cre activity (Fig. 1e, 1g). Regardless, this does not challenge the notion that U11 loss does not disrupt the basic patterning of the limb (Fig. 1b, 1g). Moreover, the difference in the phenotype observed between the mutant forelimb and hindlimb could be linked to the onset of *Prrx1*-Cre expression, as early progenitor cell depletion in the mutant forelimb may cause an insufficient number of cells to form a limb along all three axes (Supp. Fig. 2a-b). In contrast, in the mutant hindlimb, delayed progenitor cell loss may result in sufficient progenitor cells available for it to achieve patterning along all three axes (Supp. Fig. 2a-b). Nonetheless, in both the mutant forelimb and hindlimb, U11 loss causes mesomelia and reductions in bone thickness and ossification, the latter of which are observed in individuals with MODP1 (Fig. 1i, Supp. Fig. 4c-d)^21^.

The mutations in *RNU4ATAC* linked to MOPD1, RS, and LWS, and our ablation of *Rnu11*, are both expected to inhibit the minor spliceosome and disrupt minor intron splicing^11,12,14,16^. Therefore, *Rnu11* ablation can be used to glean the primary molecular defects in these diseases. Consistent with previous reports, we show that U11 loss resulted in the upregulation of minor intron retention in a subset of MIGs (Fig. 2c; Supp. Data File 1)^3,9^. Some of these MIGs are crucial for cell cycle regulation, including *Cdc45*, *Mau2*, and *Spc24*, which suggested that the limb defects observed in our mutant mice stem from cell cycle dysregulation (Fig. 2e; Supp. Data File 1)^28–30^. Indeed, we found that U11-null progenitor cells are stuck in the prometaphase to metaphase transition, which is also similar to our previous findings (Fig. 2i)^9^. Moreover, upregulation of minor intron retention in other MIGs, such as *Dna2* and *E2f6*, predicted potential S-phase defects, which was reflected in the lengthening of S-phase in the mutant forelimb and hindlimb (Fig. 3c-d; Supp. Data File 1)^31,32^. Together, these cell cycle defects reduced overall progenitor cell cycle speed (Fig. 3c-d). Consequently, we found massively elevated cell death, which was concentrated in the distal, posterior compartment of the mutant forelimb and distal, anterior compartment of the mutant hindlimb (Fig. 4c-d). This data suggested that, despite a large footprint of U11 loss in the mutant limb buds, distal progenitor cells are more susceptible to U11 loss (Fig. 4a-b, 4d). We propose that this is because these cells are the most rapidly dividing, as reported by Boehm et al.^23^, and thus do not have enough time to accommodate aberrant MIG expression. This idea agrees with our previous report of U11 loss in the developing pallium, which triggered death of the rapidly dividing radial glial cells but not the intermediate progenitors or neurons^9^. Together, the combined effect of cell cycle defects and apoptosis resulted in a severe shortage of progenitor cells that were available for limb development and would be sufficient to cause limb defects (Fig. 2g, 2i, 3c-d, 4c-d). In fact, mutations in *Cdc45* and *Dna2*, which both show elevated minor intron retention in the E11.5 mutant forelimb, are linked to primordial dwarfism in Meier-Gorlin syndrome and Seckel syndrome, respectively (Supp. Data File 1)^33,34^. These findings point to a potential role for mis-splicing of these two genes in driving the micromelia observed in individuals with MOPD1, RS, and LWS^16,17,33,34^.

Similar phenotypes of cell cycle defects and cell death were previously reported in *Dicer*^Flx/Flx^::*Prrx1*-Cre^+^ and *Ctcf*^Flx/Flx^::*Prrx1*-Cre^+^ mice^35,36^. In turn, they all show stunted limb size yet maintain skeletal organization along the proximo-distal axis, which suggests that there is a potential developmental bias for the proximo-distal axis over the antero-posterior and dorso-ventral axes. (Fig. 1f-g)^35,36^. Developmental bias, recently summarized by Wilkins^37^, suggests that a developing tissue that experiences an insult has a baseline response whereby it allocates its remaining progenitor cells to reproducibly form a basic structural unit. Moreover, these phenotypes reinforce the ideas postulated by Richardson et al.^38^, which leveraged developmental bias to argue that proximo-distal segmentation occurs first in limb development and is followed by the expansion/bifurcation of these segments. This model is borne out in our finding of a single zeugopod element and digit in the U11-null mutant forelimb rather than two zeugopod elements (Fig. 1g). Similarly, in the mutant hindlimb, which is fully patterned, the loss of the medial cuneiform and stunted digit 1 suggests that there is a critical number of cells required to form each skeletal structure (Fig. 1g; Supp. Fig. 4a). This concept of a progenitor cell threshold for skeletal element formation, first proposed by Wolpert et al.^39^, may further reinforce why the mutant forelimb, which most likely exhausts its progenitor cell population to form the basic elements of the proximo-distal axis, does not have any skeletal elements in the antero-posterior axis (Fig. 1g).

While the ideas of developmental bias can explain outcomes at the cellular and tissue level, the molecular pathways underpinning this developmental bias are unknown. Here we propose potential molecular foundations of developmental bias, as reflected in the transcriptomic changes observed in the mutant limb buds (Fig. 5a). Specifically, upregulation of 1,004 protein-coding genes in the E11.5 mutant forelimb suggested a significant transcriptomic response to cell cycle defects and cell death, which, through temporal analysis, was found to be a larger transcriptional adaptation of the limb patterning network (Fig. 5a, 5d-g). Indeed, we show through WISH that, while the developing mutant limb bud is smaller, it maintains the correct spatial organization of these signaling centers (Fig. 1d, 6a-b). Similar phenotypes were observed through x-irradiation experiments of the developing chick limb bud where, despite loss of progenitor cells, limb segment specification was maintained^40^. Together, these findings reveal potential shared mechanisms between species that maintain basic limb patterning in response to a developmental insult. In all, we show how minor spliceosome inhibition, as well as the developmental insult in the diseases MOPD1, RS, and LWS, could cause primordial dwarfism. We speculate that selection for changes in genes that could modulate the activity of the minor spliceosome might similarly result in short limbs in domesticated species.

## Acknowledgements

We thank Dr. Bo Reese from the University of Connecticut’s Center for Genome Innovation for assistance with RNAseq; Dr. Xuanhao Sun and Dr. Maritza Abril from the University of Connecticut’s Bioscience Electron Microscopy Facility for assistance with SEM imaging; Dr. Ion Mandoiu from the University of Connecticut’s Computer Science and Engineering Department for the establishment of bioinformatic platforms; and Ms. Anouk Olthof for help with bionformatic analyses. Funding for this study comes from: NIH NINDS R01NS102538 and NIH NINDS R21NS096684 to R.N.K.; NSF GRFP 2018257410 to K.D.

## Author Contributions

Conceptualization, K.D. and R.N.K.; Methodology, K.D. and R.N.K.; Investigation, K.D., C.L., A.V., G.A., and R.N.K.; Writing, K.D. and R.N.K.; Supervision, R.N.K.; Funding Acquisition, K.D. and R.N.K.

## Materials & Methods

### Animal Husbandry

Mouse husbandry and procedures were carried out in accordance with protocols approved by the University of Connecticut Institutional Animal Care and Use Committee, which operates under the guidelines of the U.S. Public Health Service Policy for laboratory animal care. The *Rnu11* conditional knockout mouse used in this study was generated and described by Baumgartner et al.,^9^. *Prrx1*-Cre was used to ablate *Rnu11* in the developing limbs^20^. The experiments described above used male and female *Rnu11*^Flx/Flx^::*Prrx1*-Cre^−^::*CAG*-loxpSTOPloxp-tdTomato^+^ (WT) mice and *Rnu11*^Flx/Flx^::*Prrx1*-Cre^+^::*CAG*-loxpSTOPloxp-*tdTomato*^+^ (mutant) mice. For embryonic harvests, E0.5 was considered noon the morning a vaginal plug was observed.

### WISH

WISH analysis was done as described previously^41^. Embryos were halved with tungsten needles (Fine Science Tools, 10130-10) after rehydration. Probes were generated by primers listed in Supp. Table 3.

### qRT-PCR

U11 loss was quantified through qRT-PCR as described in Baumgartner et al.,^9^ with normalization to *Rn7sk*. Primers used are listed in Supp. Table 3.

### SEM

SEM was performed as described previously^42^ with help from the University of Connecticut’s Bioscience Electron Microscopy Core Facility.

### Skeletal Preparation

Skeletal prep was performed as described previously^43^. Percent ossification was calculated by dividing the Alizarin red-stained region by the total long bone length.

### Bioinformatics Analysis & Data Accessibility

Total RNAseq was performed on E10.5 and E11.5 WT and mutant forelimb and hindlimb (n=3 for each sample). RNA extraction was done using TRIzol (Thermo Fisher Scientific, 15596018) per the manufacturer’s instructions. RNA sample preparation and sequencing were executed by the University of Connecticut’s Center for Genome Innovation. Library prep was done through Illumina TruSeq Stranded Total RNA Library Sample Prep Kit (RS-122-2201) with RiboZero for ribosomal RNA depletion. Sequencing was performed using Illumina NextSeq 500. Reads were mapped to mm10 using Hisat2^44^. Gene expression calls were determined through IsoEM^45^. Differential gene expression was determined using IsoDE2^45^. Minor intron retention and ORF analysis were performed as described in Baumgartner et al.,^9^. DAVID was employed for functional enrichment analysis of gene sets with significance determined by Benjamini-Hochberg adjusted *P*-value<0.05. The data discussed in this publication have been deposited in NCBI’s Gene Expression Omnibus (Drake et al., 2020) and are accessible through GEO Series accession number GSE146424.

### IF

10 um limb bud cryosections were used for IF as described previously^46^. Primary antibodies were diluted to 1:50 (MoBu-1 clone mouse anti-BrdU, Santa Cruz Biotechnology, sc-51514), 1:100 (mouse anti-Aurora-B/AIM1, BD Biosciences, 611082), or 1:500 (rabbit anti-PH3, Bethyl Laboratories, IHC-00061).

### BrdU & EdU Pulse

Timed-pregnant dams were injected with BrdU and EdU (100nmol per gram body-weight) per the paradigm described in Boehm et al.,^23^ and represented in Fig. 3a. Detection of BrdU was done through IF using the MoBu-1 clone mouse anti-BrdU antibody (Santa Cruz Biotechnology, sc-51514). Detection of EdU was performed using the Click-iT EdU Alexa Fluor 647 imaging kit (Thermo Fisher Scientific, C10340) in accordance with the manufacturer’s instructions.

### FISH

FISH for U11 was performed on 10 um limb bud cryosections as described in Baumgartner et al.,^9^.

### TUNEL

TUNEL was performed on 10 um limb bud cryosections using the *in situ* cell death detection kit, TMR Red (Roche Diagnostics), in accordance with the manufacturer’s instructions.

### Imaging & Quantification

For imaging of IF for PH3 and FISH for U11 with TUNEL, Keyence BZ-X710 was used at 20x with automatic settings for fluorescence intensity (set to the WT) and image stitching. All sections on a processed slide, which contained n=1 WT and mutant, were imaged with the same settings. To generate a heat map of TUNEL signal, all sections processed for TUNEL from a single limb bud were concatenated, stacked, and 3D projected, which was used for 3D surface plot to produce a heat map that is a 2D rendering of 3D TUNEL+ signal intensity. This was performed for a minimum of 3 limb buds per WT and mutant forelimb at E10.5 and E11.5 as well as WT and mutant hindlimb at E11.5 and E12.5. For imaging of IF for AuroraB and IF for BrdU with EdU Click-iT reaction, slides were imaged with Leica SP2 confocal microscope as described in Baumgartner et al.,^9^. Further image processing and quantification was done using IMARIS v.8.3.1 (Bitplane) and Adobe Photoshop CS4 as described in Baumgartner et al.,^9^. SEM imaging was performed using a Nova NanoSEM 450 electron nanoscope through the University of Connecticut’s Bioscience Electron Microscopy Core Facility. All representative images and traces were performed for minimum n=3, with exception for the SEM images in Fig. 1e and Supp. Fig. 2c, which were performed for n=1.

### RNA isolation and cDNA prep

Limb buds (n=3) from E10.5 and E11.5 WT and mutant embryos were dissected for RNA isolation and cDNA preparation as described by Baumgartner et al.,^9^. 100ng of RNA was used for cDNA synthesis.

### Statistical Methods

All statistical tests and results used in this manuscript are outlined in Supp. Table 4.

