## Supplementary Figures and Tables for "Minor spliceosome disruption causes limb growth defects without altering patterning"

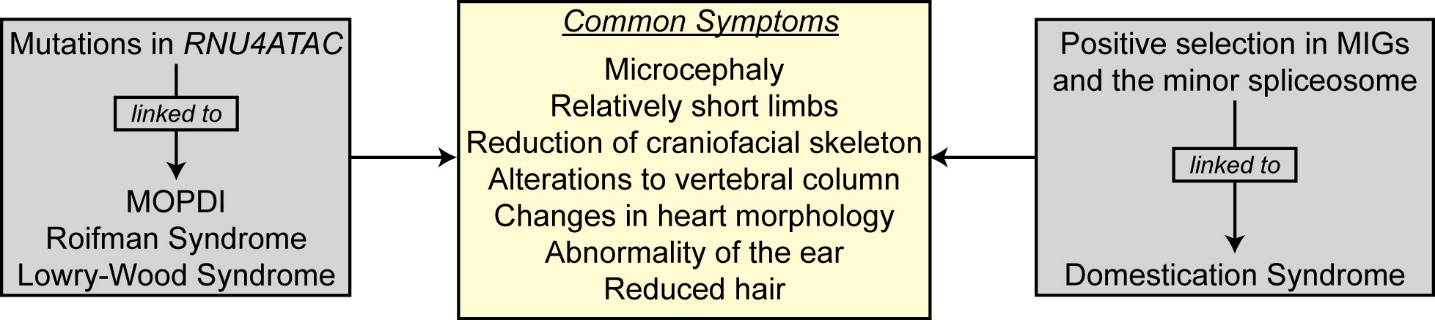

**Supplementary Figure 1 | Minor spliceosome-related diseases and domestication syndrome share overlapping symptoms.** Schematic depicting common symptoms observed in MOPD1, Roifman Syndrome, Lowry-Wood syndrome, and domestication syndrome.

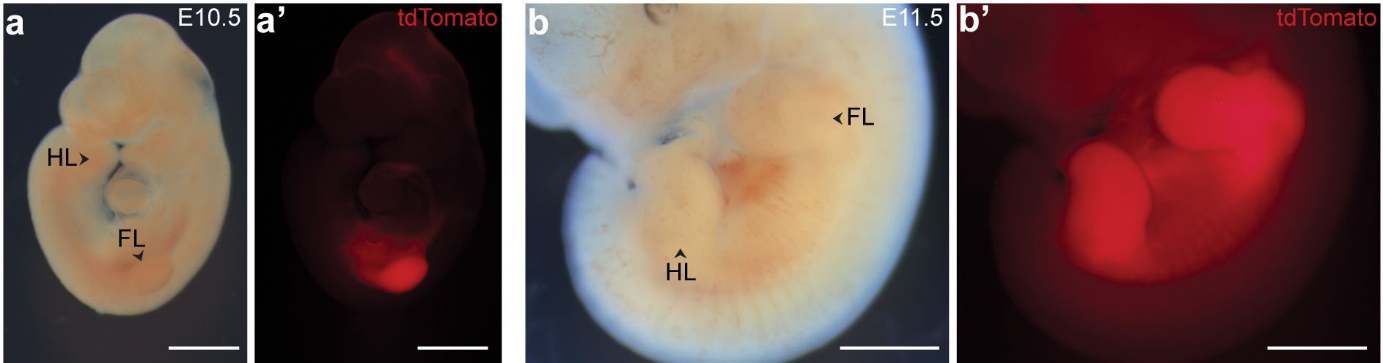

**Supplementary Figure 2 | *Prrx1*-Cre has delayed expression in the hindlimb relative to the forelimb.** **(a-b’)** Light image of mutant embryos **(a, b)** with red fluorescence imaging of tdTomato reporter **(a’, b’)** for Cre activity observed only in mutant forelimb (FL) at E10.5, FL and hindlimb (HL) at E11.5. Scale bars show 100 um.

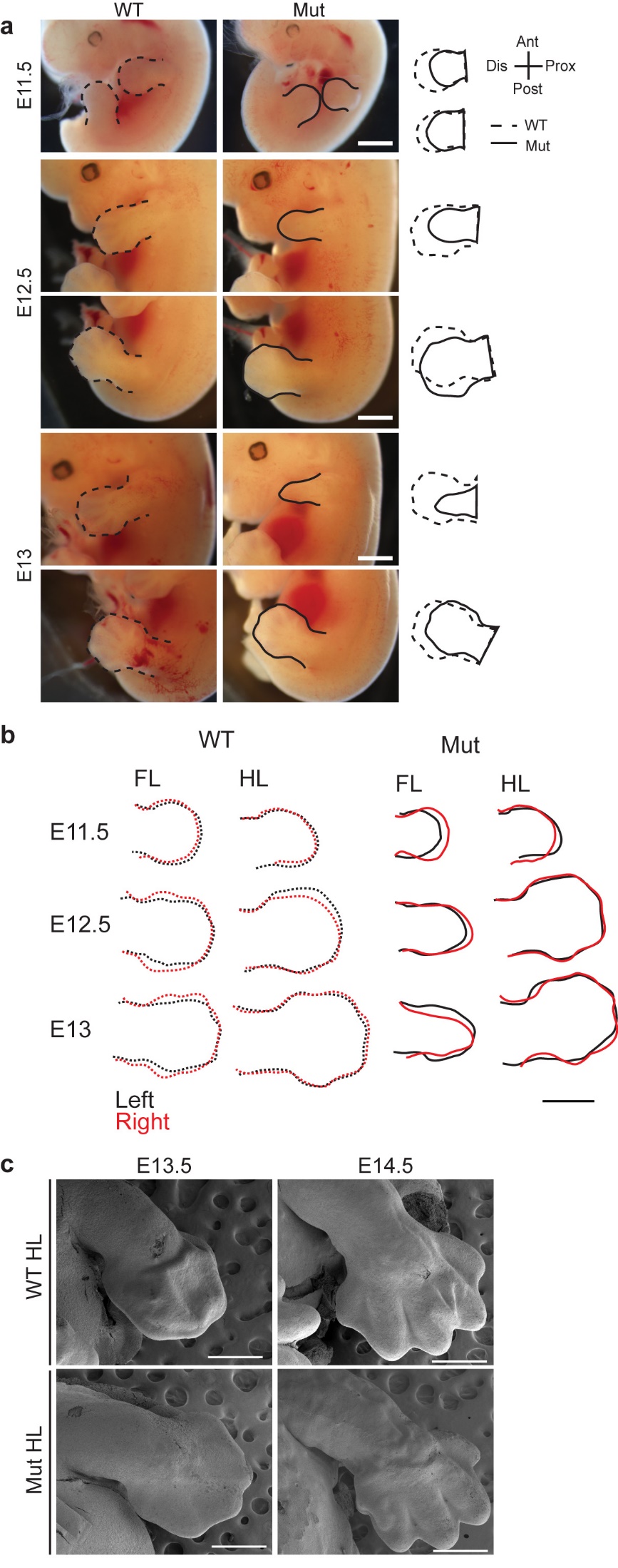

**Supplementary Figure 3 | U11 loss causes developmental defects to precipitate earlier in the forelimb versus hindlimb. (a)** Light imaged embryos at E11.5, E12.5, and E13 with traces for WT (dashed) and mutant (mut; solid) forelimb (FL) and hindlimb (HL). **(b)** Overlay of representative traces from left (black) and right (red) FL and HL buds for WT and mutant at E11.5, E12.5, and E13. **(c)** SEM of WT and mutant HL at E13.5 and E14.5. Scale bars show 100 um in **(a)** and **(b)**, 500 um in **(c)**.

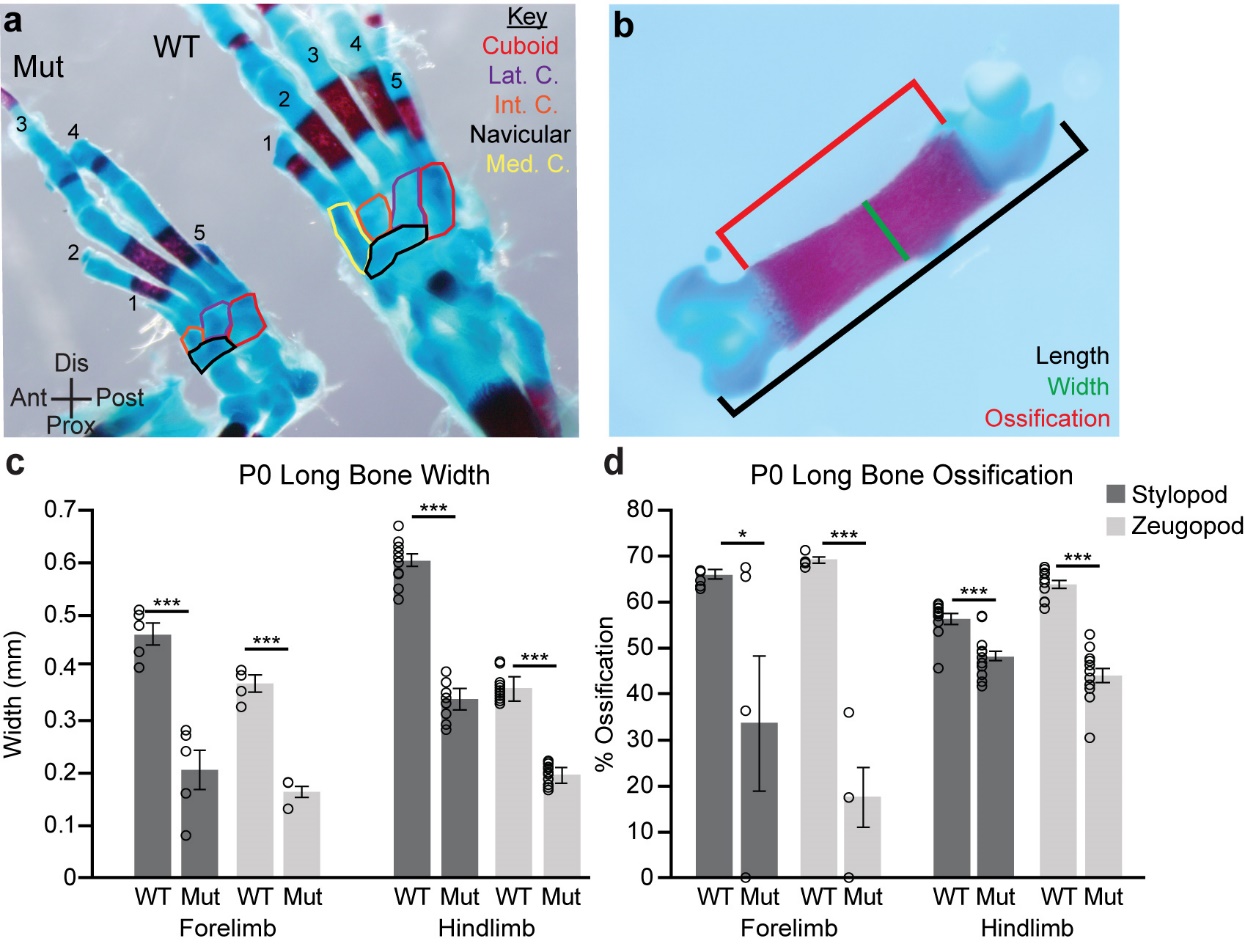

**Supplementary Figure 4 | U11 loss leads to compromised skeletal development. (a)** Skeletal preparation of mutant (mut; left) and WT (right) foot at P0 with colored traces to show metatarsals and numbers as digit labels. **(b)** Quantification schematic for long bone length, width, and ossification. **(c-d)** Bar chart showing quantification of long bone width **(c)** and ossification **(d)** for P0 WT and mutant FL and HL. Lat=lateral, Int=intermediate, Med=medial, C=cuneiform, Ant=anterior, Post=posterior, Prox=proximal, Dis=distal. Bar charts represent mean and error bars show standard error of mean. Significance determined via student’s two-tailed T-test. *=*p*<0.05, **=*p*<0.01, ***=*p*<0.001.

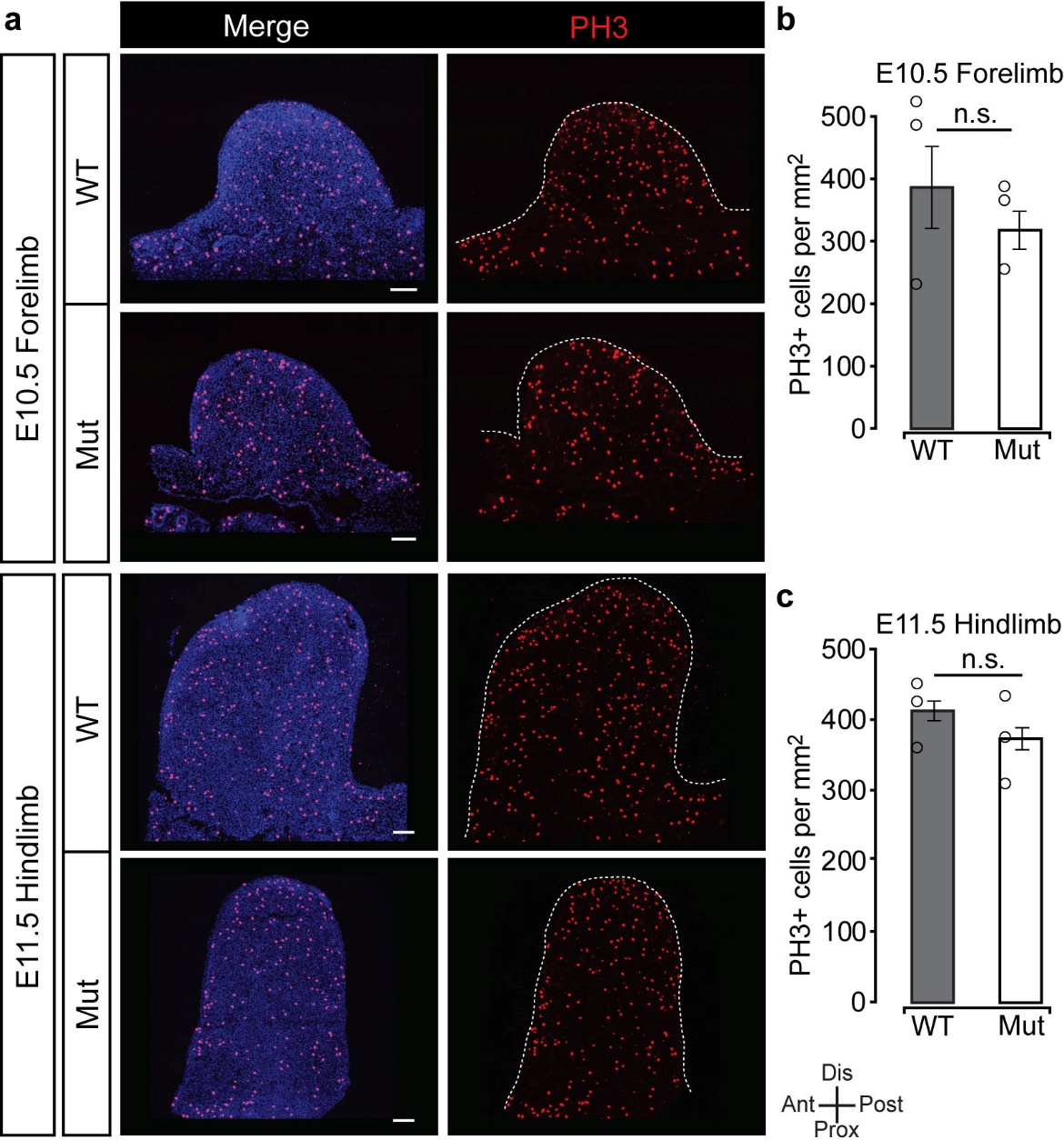

**Supplementary Figure 5 | Loss of U11 does not alter mitotic cells in the E10.5 forelimb or E11.5 hindlimb.** **(a-c)** IF for PH3 counterstained with DAPI in the E10.5 forelimb (FL) and E11.5 hindlimb (HL) for WT and mutant (mut) **(a)** with quantification **(b-c)** normalized to limb bud area. Bar charts represent mean and error bars represent standard error of the mean. Scale bars represent 100 um. Significance determined by student’s two-tailed T-test; n.s.=not significant.

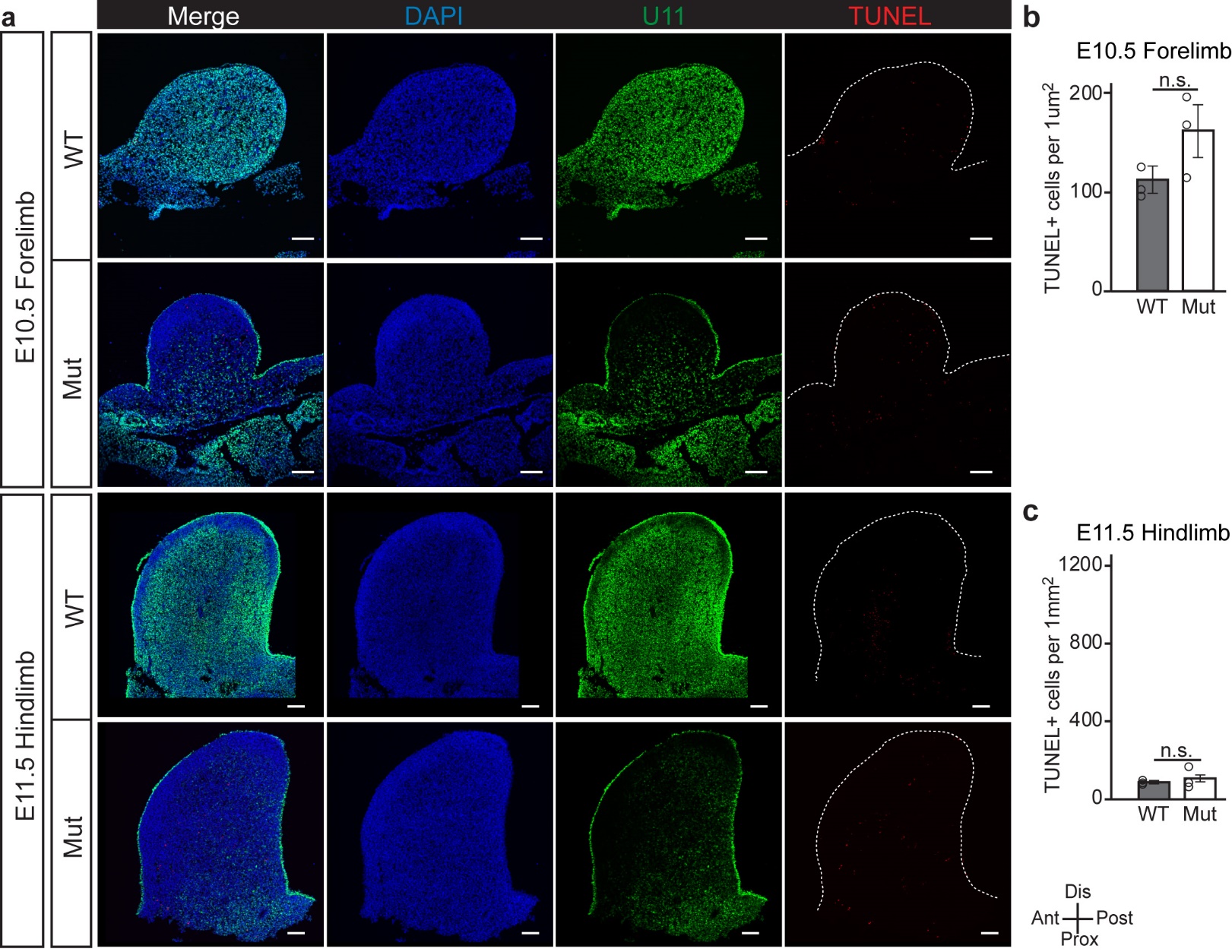

**Supplementary Figure 6 | U11 loss does not cause an increase in cell death in the E10.5 forelimb or E11.5 hindlimb.** **(a-c)** FISH for U11 with TUNEL and counterstain for DAPI in E10.5 forelimb (FL) and E11.5 hindlimb (HL) for WT and mutant (mut) **(a)** with quantification **(b-c)** normalized to limb bud area. Bar charts represent mean and error bars represent standard error of the mean. Scale bars represent 100 um. Significance determined by student’s two-tailed T-test; n.s.=not significant.

**Supplementary Tables.**

**Supplementary Table 1. Effect of minor intron retention on open reading frame (ORF).**

| Sample | # MIGs w/ incr. minor intron retention | # ORF: Extension | | # ORF: Premature Stop/Elongation | # ORF: Premature Stop/Truncation | Of Premature Stop: # NMD | Of Premature Stop: # Novel Protein |
| --- | --- | --- | --- | --- | --- | --- | --- |
| E10.5 FL | 21 | | 1 | 0 | 20 | 20 | 0 |
| E10.5 HL | 2 | | 0 | 1 | 1 | 1 | 1 |
| E11.5 FL | 134 | | 1 | 2 | 131 | 118 | 15 |
| E11.5 HL | 15 | | 0 | 0 | 15 | 14 | 1 |
| Total (unique)  # | 152 | | 2 | 3 | 147 | 134 | 16 |
| Total (unique)  % | 100% | | 1.3% | 2.0% | 96.7% | 88.1% | 10.5% |

Incr=increased, FL=forelimb, HL=hindlimb, NMD=nonsense-mediated decay

**Supplementary Table 2. Gene Ontology (GO) enrichment for differentially expressed gene sets from pairwise comparison.**

| Comparison | Differential Expr. | GO Term | # Genes | *Benjamini* |
| --- | --- | --- | --- | --- |
| E10.5 WT FL vs E10.5 Mut FL | Downregulated in E10.5 Mut FL | Transcriptional activator, RNA polymerase II transcription regulatory region sequence-specific binding | 8 | 4.72E-02 |
| E10.5 WT HL vs E10.5 Mut HL | Upregulated in E10.5 Mut HL | Extracellular region | 24 | 9.0E-03 |
| E11.5 WT HL vs E11.5 Mut HL | Downregulated in E11.5 Mut HL | Transcriptional activator activity, RNA polymerase II core promoter proximal region sequence-specific binding | 7 | 5.2E-03 |
| E11.5 WT HL vs E11.5 Mut HL | Downregulated in E11.5 Mut HL | RNA polymerase II core promoter proximal region sequence-specific binding | 8 | 5.8E-03 |
| E11.5 WT HL vs E11.5 Mut HL | Downregulated in E11.5 Mut HL | DNA binding | 15 | 6.3E-03 |
| E11.5 WT HL vs E11.5 Mut HL | Downregulated in E11.5 Mut HL | Transcription factor activity, RNA polymerase II distal enhancer sequence-specific binding | 4 | 2.2E-02 |
| E11.5 WT HL vs E11.5 Mut HL | Downregulated in E11.5 Mut HL | Proteinaceous extracellular matrix | 7 | 2.92E-02 |
| E11.5 WT HL vs E11.5 Mut HL | Downregulated in E11.5 Mut HL | Cell adhesion | 9 | 3.3E-02 |
| E11.5 WT HL vs E11.5 Mut HL | Downregulated in E11.5 Mut HL | Chromatin binding | 7 | 3.7E-02 |
| E11.5 WT HL vs E11.5 Mut HL | Downregulated in E11.5 Mut HL | Calcium ion binding | 8 | 4.4E-02 |
| E11.5 WT HL vs E11.5 Mut HL | Downregulated in E11.5 Mut HL | Transcription factor activity, sequence-specific DNA binding | 9 | 4.7E-02 |

**Supplementary Table 3. Primer sequences used for qRT-PCR and WISH probes.**

| **Primer** | **Sequence (5’ to 3’)** | **Purpose** |
| --- | --- | --- |
| *Rnu11* Forward | AAAGGGCTTCTGTCGTGAGTGGC | qRT-PCR |
| *Rnu11* Reverse | CCGGGACCAACGATCACCAG | qRT-PCR |
| *Rn7sk* Forward | CTCCAAACAAGCTCTCAAGGTCCA | qRT-PCR |
| *Rn7sk* Reverse | ATGCAGCGCCTCATTTGGATGTGT | qRT-PCR |
| *Shh* Forward | ATGCTGCTGCTGCTGGCCAGATGT | WISH |
| *Shh* Reverse | GGGCCCCGAGTCGTTGTGCGGCGC | WISH |
| *Sall1* Forward | CTCAGCTGTCAGAGCGCCTTGAAAATG | WISH |
| *Sall1* Reverse | CAGGCTATCTTGGGAAGCGTCCGC | WISH |
| *Hoxa11* Forward | GGTTCAGATCTCCGTGGTTAAG | WISH |
| *Hoxa11* Reverse | GGCTCTTAGAAGATTGCCAGAA | WISH |
| *Hoxa13* Forward | ATCCTTCAGACGCCAGCTCCTATA | WISH |
| *Hoxa13* Reverse | TGGCTGATATCCTCCTCCGTTTGT | WISH |
| *Fgf8* Forward | CAGGTCCTGGCCAACAAG | WISH |
| *Fgf8* Reverse | AGCTCCCGCTGGATTCCT | WISH |
| *Cyp26b1* Forward | TCTGCCCCTTTGCTCTTGGAAGAG | WISH |
| *Cyp26b1* Reverse | CAGGGATCCCCTTCAGCTTTTCCT | WISH |

**Supplementary Table 4. Summary of all statistical tests performed.**

| **Fig.** | **Analysis** | **n-Value** | **Mean ± SEM** | **Statistical Test** | **t/U-Value** | ***P*-Value** |
| --- | --- | --- | --- | --- | --- | --- |
| 1d | Surface Area: E11.5 FL | WT=10  Mut=6 | WT= 0.98 ± 0.04 mm^2^  Mut= 0.76 ± 0.04 mm^2^ | Student’s two-tailed T-test  (Welch’s) | 3.61533 | .002812 |
| 1d | Surface Area: E12.5 FL | WT=9  Mut=8 | WT= 2.02 ± 0.07 mm^2^  Mut= 0.97 ± 0.02 mm^2^ | Student’s two-tailed T-test  (Welch’s) | 13.15265 | 2.14313E-07 |
| 1d | Surface Area:  E11.5 HL | WT=10  Mut=7 | WT= 0.95 ± 0.04 mm^2^  Mut= 0.86 ± 0.06 mm^2^ | Student’s two-tailed T-test  (Welch’s) | 1.27939 | .110099 |
| 1d | Surface Area:  E12.5 HL | WT=9  Mut=8 | WT= 2.58 ± 0.04 mm^2^  Mut= 2.33 ± 0.06 mm^2^ | Student’s two-tailed T-test  (Welch’s) | 3.45448 | .00177 |
| 1h | Total Length:  P0 FL Stylopod | WT=5  Mut=5 | WT= 4.198 ± 0.047 mm  Mut= 1.166 ± 0.123 mm | Student’s two-tailed T-test | 22.93287 | 1.38658E-08 |
| 1h | Total Length:  P0 FL Zeugopod | WT=5  Mut=3 | WT= 3.761 ± 0.024 mm  Mut= 1.166 ± 0.123 mm | Student’s two-tailed T-test  (Welch’s) | 33.93863 | 0.000859366 |
| 1h | Total Length:  P0 HL Stylopod | WT=11  Mut=11 | WT= 3.932 ± 0.052 mm  Mut= 1.936 ± 0.044 mm | Student’s two-tailed T-test | 29.02822 | 7.99172E-18 |
| 1h | Total Length:  P0 HL Zeugopod | WT=11  Mut=11 | WT= 3.976 ± 0.070 mm  Mut= 1.438 ± 0.063 mm | Student’s two-tailed T-test | 26.85227 | 3.65808E-17 |
| 1i | Relative Length:  P0 FL Stylopod | WT=5  Mut=5 | WT= 0.501 ± 0.006%  Mut= 0.576 ± 0.214% | Student’s two-tailed T-test | -7.54062 | .000282 |
| 1i | Relative Length:  P0 FL Zeugopod | WT=5  Mut=3 | WT= 0.499 ± 0.005%  Mut= 0.424 ± 0.054% | Student’s two-tailed T-test  (Welch’s) | 4.85849 | .002827 |
| 1i | Relative Length:  P0 HL Stylopod | WT=11  Mut=11 | WT= 0.491 ± 0.003%  Mut= 0.575 ± 0.015% | Student’s two-tailed T-test | -5.53339 | 2.04295E-05 |
| 1i | Relative Length:  P0 HL Zeugopod | WT=11  Mut=11 | WT= 0.509 ± 0.003%  Mut= 0.425 ± 0.013% | Student’s two-tailed T-test | 5.70835 | .000014 |
| 2c | Median MSI:  E10.5 FL | WT=3  Mut=3 | WT= 5.537%  Mut= 10.386%  *median values | Mann-Whitney U-test | -5.226 | <0.00001 |
| 2c | Median MSI:  E10.5 HL | WT=3  Mut=3 | WT= 5.010%  Mut= 5.148%  *median values | Mann-Whitney U-test | -0.80267 | 0.42372 |
| 2c | Median MSI:  E11.5 FL | WT=3  Mut=3 | WT= 7.625%  Mut= 18.897%  *median values | Mann-Whitney U-test | -8.94649 | <0.00001 |
| 2c | Median MSI:  E11.5 HL | WT=3  Mut=3 | WT= 7.234%  Mut= 8.234%  *median values | Mann-Whitney U-test | -1.59049 | 0.11184 |
| 2g | PH3:  E11.5 FL | WT=3  Mut=3 | WT= 303.27 ± 13.42^†^  Mut= 387.29 ± 10.30^†^ | Student’s two-tailed T-test | -3.49817 | 0.024937492 |
| 2g | PH3:  E12.5 HL | WT=3  Mut=3 | WT= 330.98 ± 46.39^†^  Mut= 492.00 ± 28.59^†^ | Student’s two-tailed T-test | -3.69175 | .020986 |
| 2i | AuroraB:  E10.5 FL Prophase | WT=3  Mut=3 | WT= 9.18 ± 1.06%  Mut= 8.07 ± 1.11% | Student’s two-tailed T-test | 0.72279 | .50981 |
| 2i | AuroraB:  E10.5 FL Prometaphase | WT=3  Mut=3 | WT= 55.49 ± 0.67%  Mut= 67.44 ± 1.82% | Student’s two-tailed T-test | -6.16165 | .003521 |
| 2i | AuroraB:  E10.5 FL Metaphase | WT=3  Mut=3 | WT= 6.04 ± 0.46%  Mut= 3.95 ± 0.38% | Student’s two-tailed T-test | 3.50249 | .02484 |
| 2i | AuroraB:  E10.5 FL Anaphase | WT=3  Mut=3 | WT= 7.62 ± 1.28%  Mut= 5.31 ± 0.78% | Student’s two-tailed T-test | 1.54626 | .196942 |
| 2i | AuroraB:  E10.5 FL Telophase | WT=3  Mut=3 | WT= 21.67 ± 0.23%  Mut= 15.23 ± 2.42% | Student’s two-tailed T-test | 2.64809 | .057096 |
| 2i | AuroraB:  E11.5 HL Prophase | WT=3  Mut=3 | WT= 14.57 ± 2.44%  Mut= 8.84 ± 1.40% | Student’s two-tailed T-test | 2.03797 | .1112 |
| 2i | AuroraB:  E11.5 HL Prometaphase | WT=3  Mut=3 | WT= 58.23 ± 1.76%  Mut= 67.52 ± 3.11% | Student’s two-tailed T-test | -5.20479 | .006495 |
| 2i | AuroraB:  E11.5 HL Metaphase | WT=3  Mut=3 | WT= 5.19 ± 0.94%  Mut= 4.86 ± 0.09% | Student’s two-tailed T-test | 0.35009 | .743931 |
| 2i | AuroraB:  E11.5 HL Anaphase | WT=3  Mut=3 | WT= 6.53 ± 0.24%  Mut= 4.25 ± 0.28% | Student’s two-tailed T-test | 6.19484 | .003452 |
| 2i | AuroraB:  E11.5 HL Telophase | WT=3  Mut=3 | WT= 15.48 ± 1.48%  Mut= 14.53 ± 1.80% | Student’s two-tailed T-test | 0.40871 | .703684 |
| 3c | EdU/BrdU:  E10.5 FL Ts | WT=3  Mut=3 | WT= 9.79 ± 0.83 hrs  Mut= 13.09 ± 0.13 hrs | Student’s two-tailed T-test | -5.22469 | .006407 |
| 3c | EdU/BrdU:  E10.5 FL Tc | WT=3  Mut=3 | WT= 12.02 ± 0.71 hrs  Mut= 16.10 ± 0.30 hrs | Student’s two-tailed T-test | -6.94018 | .002264 |
| 3c | EdU/BrdU:  E11.5 HL Ts | WT=3  Mut=3 | WT= 8.65 ± 0.84 hrs  Mut= 14.68 ± 0.16 hrs | Student’s two-tailed T-test | -4.20593 | .013631 |
| 3c | EdU/BrdU:  E11.5 HL Tc | WT=3  Mut=3 | WT= 14.53 ± 0.1.31 hrs  Mut= 21.09 ± 1.59 hrs | Student’s two-tailed T-test | -4.00059 | .016122 |
| 4c | TUNEL:  E11.5 FL | WT=3  Mut=3 | WT= 50.62 ± 9.92^†^  Mut= 1195.35 ± 63.83^†^ | Student’s two-tailed T-test | -21.39637 | .000028 |
| 4c | TUNEL:  E12.5 HL | WT=3  Mut=3 | WT= 166.80 ± 23.62^†^  Mut= 901.58 ± 153.23^†^ | Student’s two-tailed T-test | -2.86248 | .045815 |
| S3c | Ossification:  P0 FL Stylopod | WT=5  Mut=5 | WT= 66.02 ± 0.98%  Mut= 33.57 ± 14.76% | Student’s two-tailed T-test | 2.53317 | .029709 |
| S3c | Ossification:  P0 FL Zeugopod | WT=5  Mut=3 | WT= 69.17 ± 0.68%  Mut= 17.53 ± 6.47% | Student’s two-tailed T-test  (Welch’s) | 6.85143 | .000476 |
| S3c | Ossification:  P0 HL Stylopod | WT=11  Mut=11 | WT= 56.34 ± 1.20%  Mut= 48.23 ± 0.95% | Student’s two-tailed T-test | 4.20736 | .000433 |
| S3c | Ossification:  P0 HL Zeugopod | WT=11  Mut=11 | WT= 63.96 ± 0.85%  Mut= 43.98 ± 1.51% | Student’s two-tailed T-test | 9.73352 | 4.97284E-09 |
| S3d | Width:  P0 FL Stylopod | WT=5  Mut=5 | WT= 0.464 ± 0.021 mm  Mut= 0.206 ± 0.015 mm | Student’s two-tailed T-test | 5.94716 | .000343 |
| S3d | Width:  P0 FL Zeugopod | WT=5  Mut=3 | WT= 0.371 ± 0.038 mm  Mut= 0.163 ± 0.011 mm | Student’s two-tailed T-test  (Welch’s) | 9.49037 | .000078 |
| S3d | Width:  P0 HL Stylopod | WT=11  Mut=11 | WT= 0.605 ± 0.013 mm  Mut= 0.339 ± 0.005 mm | Student’s two-tailed T-test | 16.08382 | 6.60316E-13 |
| S3d | Width:  P0 HL Zeugopod | WT=11  Mut=11 | WT= 0.360 ± 0.027 mm  Mut= 0.195 ± 0.015 mm | Student’s two-tailed T-test | 16.85185 | 2.7617E-13 |
| S4b | PH3:  E10.5 FL | WT=3  Mut=3 | WT=379.84 ± 64.88^†^  Mut= 312.66 ± 29.11^†^ | Student’s two-tailed T-test | 0.8296 | .453407 |
| S4b | PH3:  E11.5 HL | WT=3  Mut=3 | WT= 411.05 ± 14.30^†^  Mut= 370.67 ± 20.42^†^ | Student’s two-tailed T-test | 0.82153 | .45749 |
| S5b | TUNEL:  E10.5 FL | WT=3  Mut=3 | WT= 114.09 ± 14.33^‡^  Mut= 163.72 ± 27.15^‡^ | Student’s two-tailed T-test | -2.02333 | 0.113069 |
| S5b | TUNEL:  E11.5 HL | WT=3  Mut=3 | WT= 86.16 ± 9.97^†^  Mut= 106.86 ± 17.82^†^ | Student’s two-tailed T-test | -0.58577 | .589485 |

^†^ = cells per mm^2^ | ^‡^ = cells per um^2^ | hrs = hours | MSI = mis-splicing index
